# Human mammary 3D spheroid models uncover the role of filopodia in breaching the basement membrane to facilitate invasion

**DOI:** 10.1101/2025.08.29.672913

**Authors:** Alain Corinus, Sophie Abelanet, Julia Dubreuil, Zhenyu Zhu, Sabrina Pisano, Christelle Boscagli, Anne-Sophie Gay, Delphine Debayle, Marin Truchi, Kevin Lebrigand, Sandra Lacas-Gervais, Frédéric Brau, Xavier Descombes, Patricia Rousselle, Michel Franco, Frédéric Luton

## Abstract

Basement membranes (BM) are thin, nanoporous sheets of specialized extracellular matrix (ECM) that line epithelial tissues. They are dynamic structures that serve multiple key functions, as evidenced by numerous diseases, including cancer progression, that are associated with their alterations. Our understanding of the BM and its communication with adjoining epithelial cells remains highly fragmented due to the BM’s complex molecular architecture, the lack of molecular tools, limitations in utilizing high-resolution imaging techniques to BMs assembled on tissues, and the difficulty of assessing their functional contributions *in vivo*. Here, by combining multiple -omics analyses and advanced microscopy methodologies, we characterized the BM from two normal human mammary epithelial cell lines, MCF10 and HMLE, grown as spheroids in 3D matrices. Our findings indicate that the spheroids autonomously assemble a BM exhibiting all the molecular, structural, and biophysical characteristics of physiological BM. Using these minimalist model systems, we show for the first time that the laminins, perlecan, and the hemidesmosomes are all arranged along a shared porous lattice defined by the collagen IV molecular network. Next, we demonstrate that the invasion-promoting *PSD4*/EFA6B knockout, found in patients with breast cancer, decreases the expression of BM components and their assembly on the spheroid surface. We then show that invasive spheroids develop enlarged pores in the BM via filopodia-like plasma membrane extensions, which further expand in a protease-dependent manner, thereby facilitating the passage of invasive cells.

**Significance Statement:** The basement membrane (BM) directs epithelial tissue architecture and behavior. How tumor cells breach this barrier during invasion remains poorly understood. Using minimalist 3D spheroid models mimicking physiological BM assembly, we reveal a novel infiltration mechanism. Filopodia-like protrusions perforate and widen BM pores. This facilitates cell dissemination through sequential protease-independent and -dependent steps.

## Introduction

Despite significant advances in treatment, breast cancer (BC) remains a leading cause of death by cancer among women worldwide. The vast majority of breast tumors are of ductal origin, among which 20% are diagnosed *in situ* (DCIS, Ductal Carcinoma *In Situ*), meaning they have not crossed the basement membrane (BM). At least one third of these tumors remain *in situ* and benign, while the others progress to an invasive stage (IDC, Invasive Ductal Carcinoma). No predictive markers nor specific treatments exist to predict and prevent the critical transition of DCIS to IDC. Thus, patients with *in situ* tumors are treated as if they had malignant tumors, causing unnecessary human hardship and economic cost. It is therefore crucial to elucidate the mechanisms underlying BM infiltration and their regulatory pathways to develop novel therapeutic strategies (1, 2).

The transition from DCIS to IDC corresponds to the infiltration by tumor cells of the basement membrane (BM), a specialized layer of extracellular matrix surrounding epithelial tissues. The BM is a thin microporous meshwork that contributes to the organization of the breast tissue, it is a reservoir of cytokines and represents the first physical barrier to the dissemination of tumor cells into the stroma (3). The BM consists of two superimposed protein networks: type IV collagen facing the stroma and laminins facing the cells. The two networks are connected via cross-linking proteins among which the best described are the nidogen and perlecan (4, 5). Laminins are trimers formed in the Golgi apparatus, secreted and then assembled at the cell surface by receptors, mainly integrins. The cellular origin of the components of the BM is disputed and may vary among tissues. However, the breast epithelium which comprises two cell types is able to produce autonomously all laminins and major components of the BM (6–8). Despite its recognized function as a barrier controlling tumor dissemination, the mechanisms used by invasive cells to breach the BM remain to be determined.

It has been proposed that the crossing of the BM could be mediated by enzymatic digestion or by the application of physical forces to create openings (9–12). Recent studies suggested that by modifying the expression of component(s) of the BM, thus changing its composition and structural organization, tumor cells would produce an altered BM permissive to infiltration (13, 14). In support of this hypothesis, the DCIS-IDC transition in patients is accompanied by a change in the expression of genes encoding for extracellular matrix proteins and their receptors (15, 16). How these distinct mechanisms function individually at the molecular level, how they are temporally coordinated and whether regulatory relationships link them remain to be elucidated.

We reported that in BC patients the reduction of EFA6B (Exchange factor for Arf6) expression is associated with metastases and reduced survival. Further, the knock-out (KO) of EFA6B in human mammary cells facilitates the DCIS-IDC transition in a xenograft model and induces cell invasion *in vitro* in 3D collagen type I (17, 18). The transcriptomic analyses showed that the ontologies associated with EFA6B KO cells are the same as those associated with the DCIS-IDC transition *in vivo*, and that major signatures are related to ECM remodelling (18). All these results, prompted us to study BM infiltration in human mammary invasive cell models.

In this study, we validated a minimalist 3D model system to investigate the architectural organization of the BM surrounding human mammary cell spheroids and demonstrated the role of filopodia in breaching the BM. We report that human mammary spheroids autonomously assemble a BM resembling physiological BMs, in which hemidesmosomes and perlecan colocalize with superimposed laminin and collagen IV networks that delineate micropores. Three-dimensional confocal fluorescence reconstruction and electron microscopy reveal that these BM micropores are traversed by small, dynamic, F-actin–rich filopodia. In invasive spheroids, the BM is severely altered, displaying enlarged pores generated by stable elongated filopodia that precede the protease-dependent cell infiltration. Our data support a model in which BM infiltration by invasive cells involving compositional changes, filopodia-driven perforation, and protease-mediated pore expansion.

## Results

### BM components repertoire of MCF10 and HMLE cell lines

We first assessed whether the normal human mammary cell line models MCF10 and HMLE express a repertoire of BM components similar to that of the human mammary gland. We compared their gene expression profiles using publicly available single-cell RNA-seq (scRNA-seq) datasets of both *in vitro* models (19, 20) and the human breast cell atlas (21). First, using a previously defined 184-gene BM signature (22) we performed a correlation analysis which indicated that the MCF10 and HMLE cell lines display a BM signature similar to that of the epithelial compartment of the human mammary gland (**Fig. 1A**). Second, we focused on the expression of the genes encoding for the core components, that is the laminins (*LAM*) and collagen IV (*COL4A*) individual chains as well as the cross-linking proteins nidogen (*NID*) and perlecan (*HSPG2*). We found that both cell lines exhibit a similar expression profile, which closely resembles that of the epithelial compartment of the human mammary gland. There is a strong expression of the genes encoding the LAM chains that form the LAM332, LAM511 and LAM521 as previously described by immunohistochemistry. Although, little *LAMA1* is detected, LAM111 expression has been reported in the mammary gland (23–25). Consistent with the epithelial compartment of the human mammary gland, MCF10 and HMLE express high levels of genes encoding the ubiquitous collagen IV chains which assemble the main trimer α1α1α2 and to a lesser extent the α5α5α6 trimers. While *HSPG2* is clearly detected in all samples, *NID1* and *NID2* are mostly detected in HMLE, less in MCF10 cells or the epithelial compartment of the human mammary gland (**Fig. 1B**).

**Figure 1:**
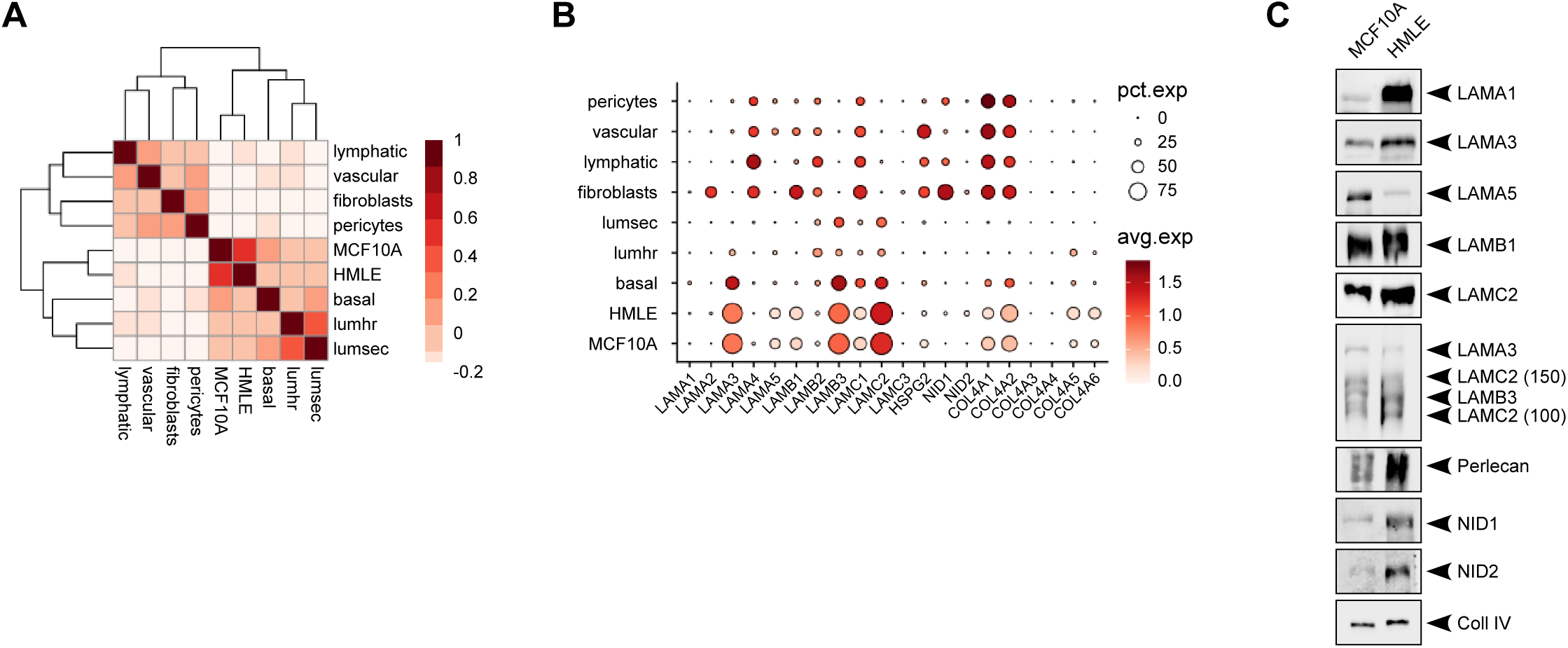
MCF10 and HMLE cell lines express a similar repertoire of BM components to that of the human mammary gland. **A**) Heatmap of Pearson’s correlation coefficient between average scRNA-seq expression profiles of 184-gene BM signature(22) in human mammary cell lines and cell types from the human mammary gland. **B**) Percentage and average level of expression of the structural core components of the BM in MCF10 and HMLE cell lines, and in cell types of the human mammary gland. **C**) Proteins from MCF10 and HMLE culture supernatants were isolated and the expression of the indicated secreted proteins analyzed by immunoblot.

Next, we refined the scRNA-seq data by conducting immunoblots for the major BM components synthesized (cell lysate, **Fig. S1**) and secreted (culture medium, **Fig. 1C)** by the MCF10 and HMLE cells. We found that both cell lines secreted similar levels of LAM332 and Coll IV α1 chain. MCF10 secreted more LAMα5 while HMLE more LAMα1 suggesting more LAM111 trimers in HMLE BM and more LAM511/521 trimers in MCF10 BM. Regarding the cross-linkers, MCF10 cells predominantly secreted perlecan while HMLE cells expressed in addition significant amounts of both nidogen 1 and 2 as revealed by scRNA-seq (**Fig. 1A,B**). Altogether, these results show that MCF10 and HMLE, despite some differences, closely mirror the expression profile of BM components found in the human mammary gland. They also indicated that these cell lines autonomously produced and secreted all the core components necessary for BM assembly.

### BM assembled on the surface of MCF10 and HMLE spheroids

Next, we investigated whether the secreted components assembled into a BM on the surface of spheroids grown in 3D Matrigel. Matrigel which was chosen specifically because it inhibits invasion and preserves an intact BM even in invasive spheroids (26). The time-course assembly of Coll IV α1 revealed that the BM was fully assembled at 72h-96h (**Fig. S2A**), therefore all subsequent experiments were conducted at 96h unless otherwise indicated. As shown in **Fig. 2A**, we found that MCF10 and HMLE spheroids assembled a BM containing the LAMα1, LAMα5, LAMγ1 and LAMγ2, the cross-linker proteins perlecan and nidogen 1, and the collagen IV. Because Matrigel is derived from murine cells and contains some BM components that might help assemble the BM, we stained the MCF10 spheroids with murine-specific antibodies directed against nidogen 1 or LAMβ1. We observed a general background reflecting the staining of the Matrigel itself but neither chains were found deposited at the surface of the spheroids (**Fig. S2B,C**). Further, spheroids grown in fibrillar type I collagen also assembled a BM (**Fig. S2D**). These observations suggest that the cells neither use nor need Matrigel components to assemble their own BM.

**Figure 2:**
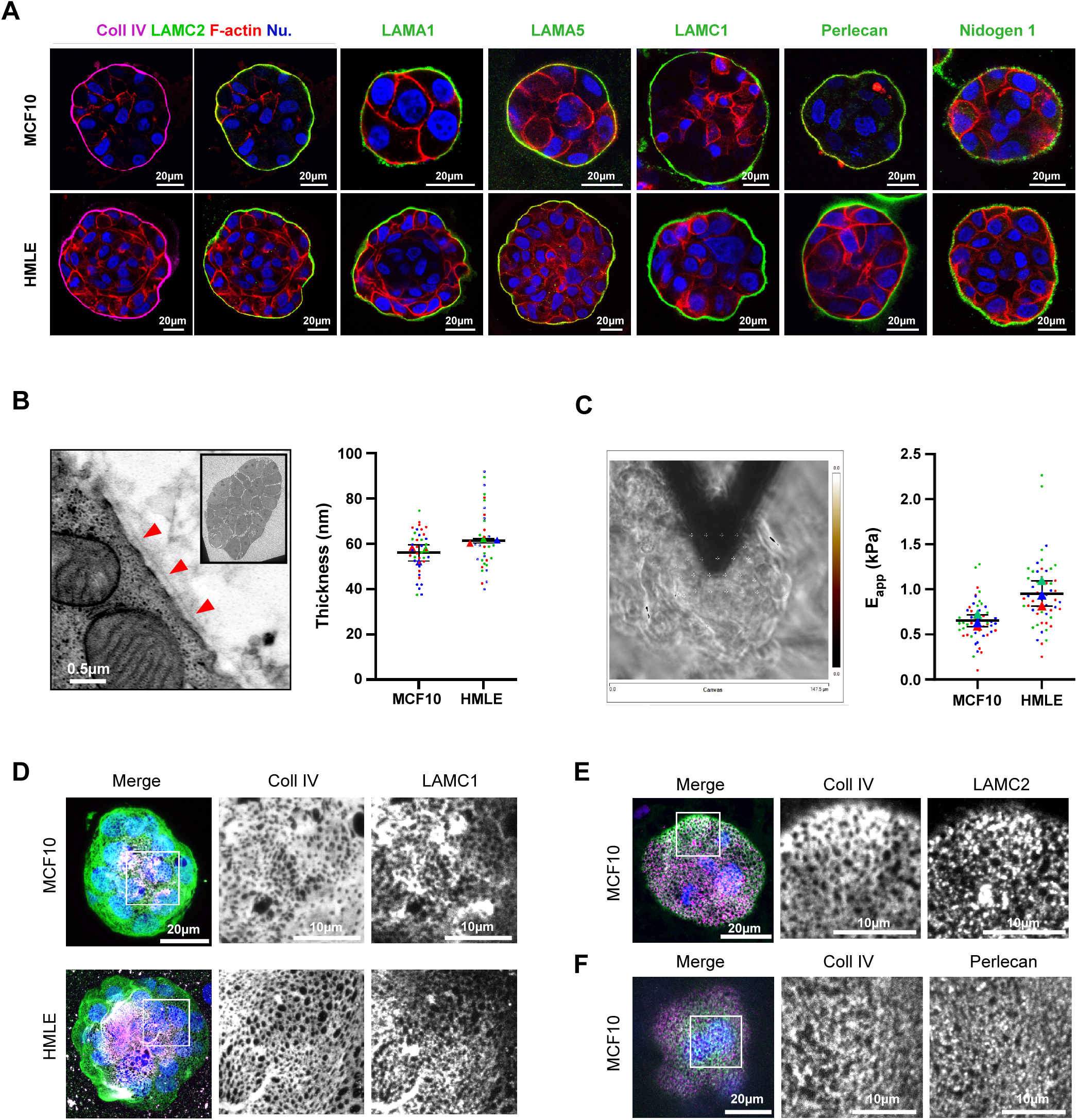
MCF10 and HMLE cell spheroids grown *in vitro* assemble a BM similar to that of previously reported physiological BM. **A**) Representative confocal immunofluorescence images of the indicated spheroids grown 96h in Matrigel and stained for the indicated BM components. **B**) TEM image of the surface of an MCF10 spheroid (inset: whole spheroid). Red arrowheads point to the thin dark line formed by the BM lining the spheroid. The right graph reports the quantification of the BM thickness of MCF10 spheroids (n=20) and HMLE spheroids (n=29), N=4 independent experiments. **C**) Phase contrast image showing the probe sampling the BM on top of a MCF10 spheroid. The right graph reports the stiffness measurement of the BM lining MCF10 spheroids (n=55) and HMLE spheroids (n=54). The colour code allows to discriminate the individual data sets corresponding to the N=3 independent experiments. **D**) MIP of confocal immunofluorescence serial images of MCF10 (top) and HMLE (bottom) spheroids grown 96h in Matrigel and stained for collagen IV (green), LAMC1 (magenta), F-actin (red) and nuclei (blue). Left panels show representative merged MIP images of the bottom third of the spheroids. Right panels show zoom-in images of the area marked in the left panels for Coll IV and LAMC1. **E**) As described in D), MIP of the bottom third of a MCF10 spheroid stained for Coll IV (green) and LAMC2 (magenta). **F**) As described in D), MIP of the bottom third of a MCF10 spheroid stained for Coll IV (green) and Perlecan (magenta).

The ultrastructure of the BM assembled on the surface of MCF10 spheroids was resolved using scanning electron microscopy (SEM). We observed a nanoporous meshwork resembling that of BM extracted from *in vivo* tissues (27, 28) (**Fig. S2E**). We also examined the organization of the BM and measured its thickness by transmission electron microscopy (TEM). In both cell models, we observed the correct positioning of the lamina lucida (laminin network) and lamina densa (collagen IV network) facing the cells and the Matrigel, respectively. The thickness of both MCF10 and HMLE BMs (≈60nm) is comparable to that of the BM measured *ex vivo*, with a thicker BM in HMLE compared to MCF10 due to a thicker lamina densa, suggesting a more multilayered organization of the collagen IV network (28, 29). (**Fig. 2B et Fig. S2F**). Finally, by atomic force microscopy (AFM) we assessed the stiffness of both MCF10 and HMLE spheroid surfaces on which the BM is assembled, and again we found values (≈0.5-1kPa) similar to that of *ex vivo* BMs (27–30) (**Fig. 2C**). We concluded that MCF10 and HMLE spheroids autonomously assemble BMs with composition, organization, and biophysical properties similar to those of the human mammary gland or other epithelial tissues. Therefore, they represent pertinent model systems for studying the infiltration of an endogenously assembled BM by invasive cells.

Before exploring how invasive cells traverse the BM, we aimed to determine its architectural organization by examining the relative positioning of the LAM and Coll IV networks, the hemidesmosomes (HD) and the perlecan. Using immunofluorescence and confocal serial Z-sectioning, spheroids stained for Coll IV and LAMγ1 (a chain common to the network-forming laminins (111, 511 and 521) demonstrated identical and superimposed meshworks (**Fig. 2D**) with pores of about 0.6µm^2^ (**Fig. 3D,4E**) which corresponds to the surface area of the polygones formed by the collagen IV molecules. Given that the porosity of laminin networks, based on molecular size, is approximately 100 times smaller than that of collagen IV, we propose that the collagen IV meshwork provides a structural scaffold that organizes the spatial organization of the laminin network and the overall porosity of the BM (31, 32). We labelled the HD with LAMγ2 specific antibodies. Though the staining appeared more dot-like as expected for HD puncta, we found that they also colocalized all along the Coll IV network, (**Fig .2E**). Finally, perlecan was found to form a quasi continuous meshwork overlapping with that of the Coll IV (**Fig. 2F**). Thus, our data indicate that the main constituents of the BM are all arranged along a shared porous lattice defined by the Coll IV meshwork.

**Figure 3:**
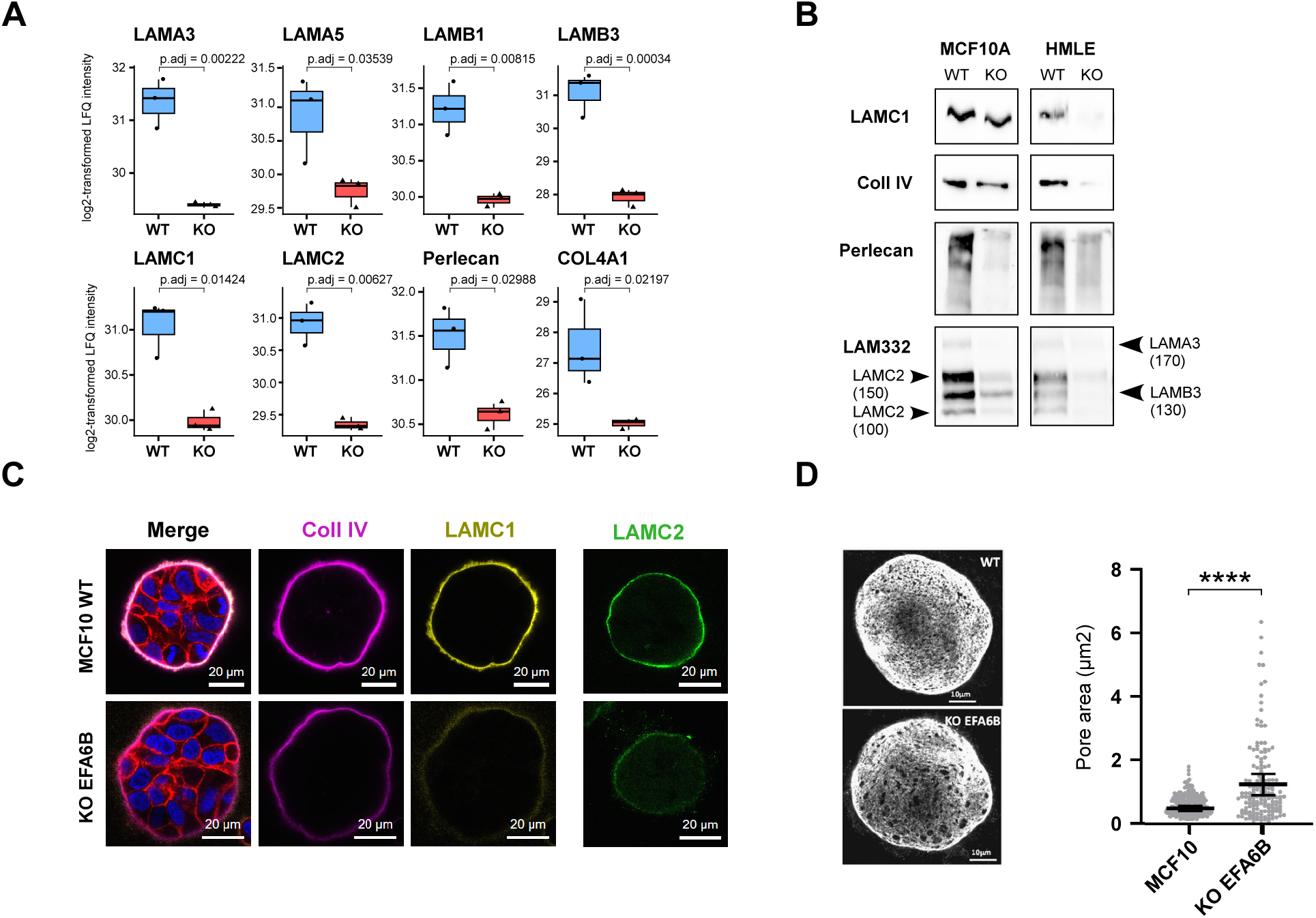
**A**) Log2-transformed LFQ intensities of BM components detected by mass spectrometry in the supernatant of MCF10WT and MCF10 EFA6B KO cells culture medium. **B**) Expression levels of the core components of the BM secreted by MCF10 and HMLE WT cell lines and a corresponding EFA6B KO clone. **C**) Representative confocal immunofluorescence images of MCF10 WT (top) and EFA6B KO (bottom) spheroids grown 96h in Matrigel and co-stained for Coll IV (magenta), LAMC1 (yellow), F-actin (red), nuclei (blue), or stained for LAMC2 (green). **D**) MIP of confocal immunofluorescence serial images of MCF10 WT (top) and EFA6B KO (bottom) spheroids grown 96h in Matrigel and stained for collagen IV. The right graph reports the surface area of each individual pore (n) of the BM of the indicated cell spheroids from three independent experiments calculated using a custom-made software (see Materials and Methods). MCF10WT mean±SEM = 0.605±0.022 (n=210); EFA6B KO mean±SEM = 1.272±0.101 (n=149)

**Figure 4:**
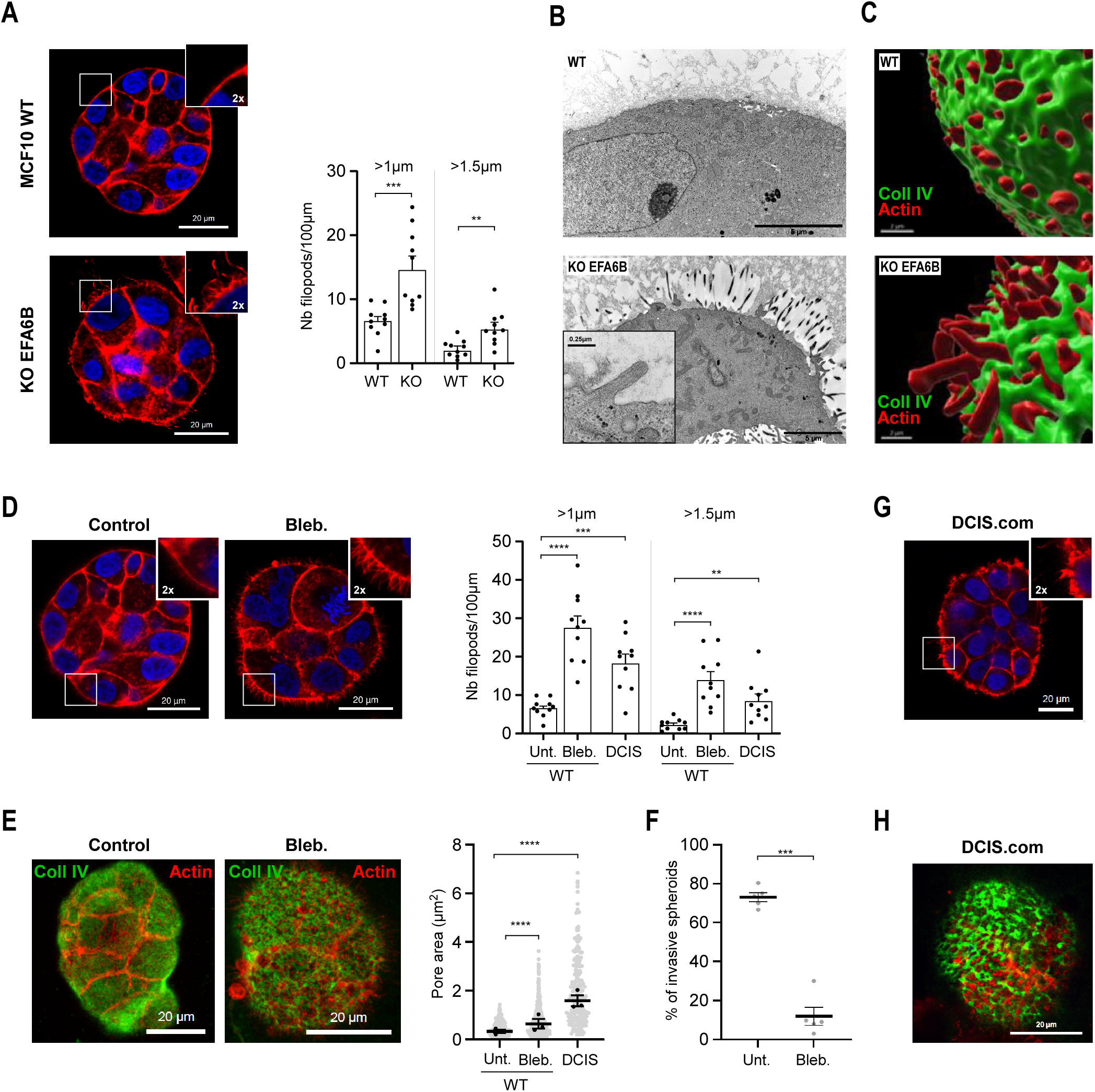
**A**) Representative confocal immunofluorescence images of MCF10 WT (top) and EFA6B KO (bottom) spheroids grown 96h in Matrigel and stained for F-actin (red), nuclei (blue). Inset: 2X zoom of the indicated area. The graph reports the number of filopodia per 100µm of the spheroids’ circumference of length over 1µm (MCF10WT mean±SEM = 6.54±0.73; EFA6B KO mean±SEM = 14.77±1.86) or over 1.5µm (MCF10WT mean±SEM = 2.20±0.42; EFA6B KO mean±SEM = 5.49±0.85). n=10, N=3. *P* values calculated using an unpaired t test with Prism software. \*\**p*<0.01; \*\*\**p*<0.005. **B**) Representative TEM images of MCF10 WT (top) and EFA6B KO (bottom) cells at the periphery of the spheroids. The inset is a zoom-in magnified image of a filopodia revealing the presence of F-actin bundles oriented lengthwise. **C**) 3D reconstruction of the collagen IV (green) and F-actin (red) staining of MCF10 WT (top) and EFA6B KO (bottom) spheroids. The images were obtained after deconvolution and 3D reconstruction using the Huygens and Imaris softwares, respectively. **D**) Representative confocal immunofluorescence images of untreated (left) and blebbistatin-treated (right) MCF10WT spheroids grown 96h in Matrigel and stained for F-actin (red), nuclei (blue). Inset: 2X zoom of the indicated area. The graph reports the number of filopodia per 100µm of the spheroids’ circumference of length over 1µm (WT untreated (Unt.): mean±SEM = 6.54±0.73; WT+Bleb.: mean±SEM = 27.39±2.86; DCIS.com: mean±SEM = 18.21±2.26) or over 1.5µm (WT intreated (Unt.): mean±SEM = 2.20±0.41; WT+Bleb.: mean±SEM = 13.77±2.12; DCIS.com: mean±SEM = 8.46±1.70). n=10, N=3. *P* values calculated using an unpaired t test with Prism sofwtare. \*\**p*<0.01; \*\*\**p*<0.005; \*\*\*\**p*<0.0001. **E**) MIP of confocal immunofluorescence serial images of untreated (left) and blebbistatin-treated (right) MCF10WT spheroids grown 96h in Matrigel stained for collagen IV (green), F-actin (red). The right graph reports the surface area of each individual pore of the BM of the indicated cell spheroids from three independent experiments calculated using a custom-made software: WT untreated (Unt.) n=201, mean±SEM = 0.474±0.019; WT+Bleb. 50µM treated MCF10WT for the last 16h n=559, mean±SEM = 0.718±0.025; DCIS.com. n=202, mean±SEM = 2.017±0.101. **F**) Quantification of the percentage of cell aggregates (n =100) with invasive protrusions of EFA6B KO cell spheroids grown in collagen for 3 days and treated or not (Unt.) with Blebbistatin 50µM for the last 16h (Bleb.). N = 3, average ± SEM. *P* value calculated using a paired t test, \*\*\**p*<0.001. **G**) Representative confocal immunofluorescence image of DCIS.com spheroids grown 96h in Matrigel and stained for F-actin (red), nuclei (blue). Inset: 2X zoom of the indicated area. In E) The graph reports the number of filopodia per 100µm of the spheroids’ circumference of length over 1µm or over 1.5µm. n=10, N=3. *P* values calculated using an unpaired t test with Prism sofwtare. \*\*\*\**p*<0.0001. **H**) MIP of confocal immunofluorescence serial images of DCIS.com spheroids grown 96h in Matrigel stained for collagen IV (green), F-actin (red).

### BM of EFA6B KO invasive spheroids

To investigate the mechanisms of BM infiltration, we used EFA6B KO human mammary cells as a model of collective invasion in 3D culture. We previously reported that reduced expression of EFA6B in tumor is associated with invasive BC in both animal model and patients. Gene expression profiling of EFA6B KO cells revealed enrichment of ECM–related gene ontologies (GO), mirroring patterns observed IDC human tumor samples (15, 16, 18). To further examine whether the BM of EFA6B KO cells is altered, thereby facilitating invasion, we first analyzed the secretome by mass spectrometry. The proteomic analysis showed that the levels of expression of many BM core components were down-regulated in MCF10 EFA6B KO cells (**Fig. 3A and Fig. S3A**) and GSEA revealed that several GO related to ECM organization, including the BM, and its cell-interaction were affected (**Fig. S3B**). Immunoblot analyses confirmed that invasive EFA6B KO MCF10 and HMLE cells secreted lower levels of the BM core components (**Fig. 3B**). Consequently, we observed a reduced accumulation of BM components at the surface of the KO cell aggregates in both MCF10 and HMLE models (**Fig. 3C, Fig. S3C**).

Since a decrease of BM components could alter its organization, we next analysed its architecture by optical sectioning of spheroids stained for Coll IV using laser confocal microscopy. Maximum intensity projection (MIP) of the bottom regions of the spheroids revealed a distinct increase of the surface area of the pores. The quantification of the surface area using a custom-made software showed that in average the pore size of the WT aggregates BM was 0.6µm^2^ while that of the EFA6B KO spheroids was doubled to 1.27µm^2^. Notably, in EFA6B KO spheroids we detected a fraction of pores greater than 2µm^2^, which were absent in WT spheroids (**Fig. 3D**). Thus, EFA6B KO induced down-regulation of the expression of BM core components and led to a significant increase in the pore size of the BM meshwork.

### Filopodia-driven pore enlargement facilitates BM cell infiltration

Confocal microscopy revealed that EFA6B KO spheroids displayed longer F-actin-enriched filopodia-like plasma membrane protrusions (hereafter named filopodia for simplicity) than WT spheroids (**Fig. 4A**). The frequency distribution of filopodia length showed that most are below 0.5µm, but the EFA6B KO spheroids display a larger proportion of longer filopodia, greater than 1µm (**Fig. 4A**). The long filopodia at the surface of the EFA6B KO spheroids were readily visible by TEM (**Fig. 4B**). High magnification confirmed that these filopodia are supported by parallel F-actin bundles (**Fig. 4B, inset**). The 3D reconstruction of spheroids labelled for the collagen IV and F-actin showed that these filopodia traversed the BM pores (**Fig. 4C**, **Fig. 3D**).

Because the EFA6B KO invasive cells display long filopodia and large pores in their BM, we postulated that the filopodia could pass through the nanoporous BM sheet to generate larger pores. To test this hypothesis, we first induced filopodia formation and examined the impact on the pore size. Exposing the MCF10WT spheroids to blebbistatin triggered the extension of many filopodia (13) (**Fig. 4D**), which spatially coincided with enlarged pores in the BM (**Fig. 4E**), establishing a causal relationship between filopodia formation and pore enlargement. Yet, blebbistatin inhibited invasion in collagen suggesting that actomyosin contractility is subsequently required for cell dissemination (**Fig. 4F**). Second, we studied the invasive MCF10 derived tumor cell line DCIS.com that was previously shown in 2D culture to present spontaneously more filopodia than MCF10WT (33). We observed that DCIS.com cell spheroids were also extending more protrusions than the MCF10WT (**Fig. 4D,G**). Strikingly, the pores of the BM of the DCIS.com spheroids were commensurately much larger than those of the WT (**Fig. 4E,H**). These results indicate that membrane protrusions extended by invasive cells perforate the BM and shape enlarged pores.

### Live-imaging analysis of filopodia and cell invasion

Next, we hypothesized that filopodia could facilitate invasive cells in traversing the BM barrier by enlarging pores. To test this, we analyzed the temporal dynamics of filopodia using phase-contrast videomicroscopy. Spheroids embedded in collagen I gels were monitored over 72h. To image entire spheroids, 20 cross-sectional images (10 µm steps) were acquired every 30 minutes. Custom-made software was developed to automatically detect plasma membrane extensions and extract key parameters, with the exception of width which was measured manually due to technical limitations (see Methods). Similar to spheroids in Matrigel (**Fig. 4**), we measured a greater number of protrusions in invasive EFA6B KO spheroids compared to WT spheroids in collagen I. This was accompanied by increased protrusion length and width, and notably, a markedly longer lifespan, indicating that protrusions generated by invasive cells were overall more stable (**Fig. 5A-E and Movie S1**). Frequency distribution analysis revealed that the vast majority of the filopodia had a lifespan of less that 30min. In WT spheroids, the number of filopodia with a lifespan greater than 30min steadily declined, with 95% displaying a lifespan below 90min (**Fig. 5F and Fig. S4A**). In contrast, in KO spheroids only 85% had a lifespan below 90min, and a distinct population of highly stable filopodia, absent in WT spheroids, emerged at 360min (**Fig. 5F**). Considering that only one to three invasion events were observed per aggregate, this limited number of stable filopodia could be sufficient to support the rare occurrence of invasion. This observation is consistent with the detection of a small fraction of very large pores in the BM of EFA6B KO spheroids (**Fig. 3D**). Cell invasion involves protease-dependent and -independent mechanisms. Thus, we examined the impact of the broad-spectrum protease inhibitor GM6001 on membrane protrusions characteristics. First, we confirmed that GM6001 treatment efficiently inhibited MCF10 EFA6B KO invasion in collagen I **(Fig. S4B).** We then found that GM6001 induced the formation of more numerous and longer, yet thin filopodia that never led to cell invasion (**Fig. 5A-E and Movie S1**). Despite these differences, the lifespan remained similarly high to that observed in invasive EFA6B KO spheroids, and frequency distribution analysis revealed the presence of the small population of long-lived filopodia at 360min. It suggests that protease activity is not required for the extension of stable pro-invasive membrane protrusions, but rather for the subsequent filopodia and pore enlargement necessary for crossing the BM. As indicated by the solidity index (ratio of pore area to convex envelope area), which captures the curvature of protrusive extensions, spheroids treated with GM6001 tend to project less linear filopodia, potentially reflecting an adaptive search for paths through the collagen fiber matrix (**Fig. 5G**). Next, we investigated the steps that follow the formation of stable filopodia that leads to subsequent cell invasion. To this end, we performed phase contrast and fluorescence microscopy to simultaneously track filopodia and nuclei stained with Hoechst 33342 (**Fig. 5H, Fig. S4C-F, and Movies S2-S6**). We observed that nuclei underwent deformation, and in some cases partial fragmentation, to enter the filopodia prior to cell elongation away from the aggregate, eventually followed by additional cells. To quantify this process, we measured filopodia thickness at three key time points: once the nascent filopodium stabilized, just before nuclear entry, and after the nucleus had clearly entered the filopodium (**Fig. 5I**). Our observations revealed a phase of filopodia enlargement preceding nuclear translocation. Once the filopodia thickness reached a threshold the nucleus moved in further increasing the width, followed by overall cell dissemination. Altogether, these findings support a model in which EFA6B KO invasive cells first extend long-lived filopodia, followed by a protease-dependent enlargement phase that facilitates cellular invasion, likely by widening existing BM pores.

**Figure 5:**
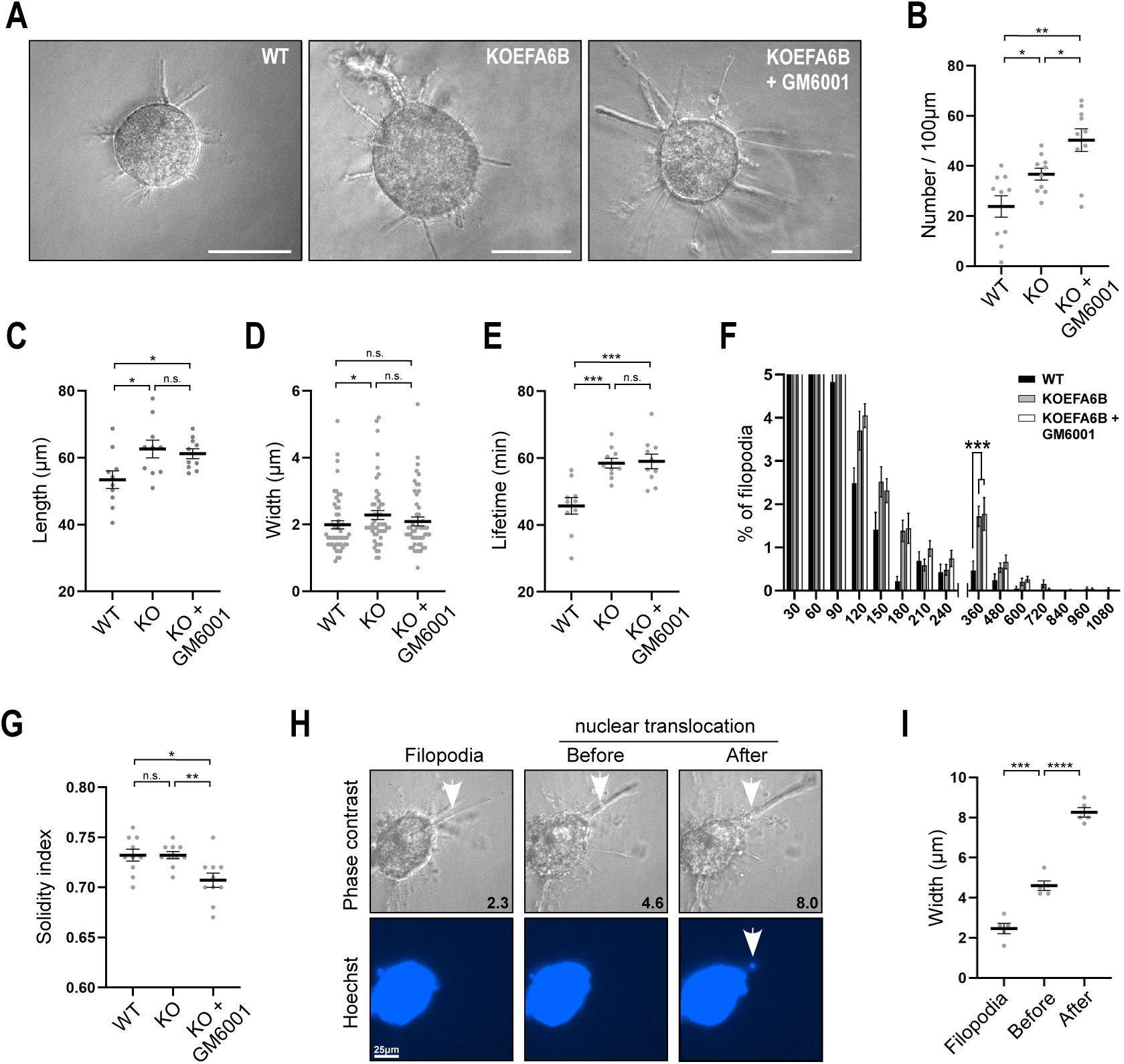
**A**) Representative images of phase contrast videomicroscopy of MCF10 WT (left), EFA6B KO (middle) and GM6001-treated EFA6B KO (right) spheroids grown in collagen for 72h. Scale bar 100µm. **B, C, D, E, G**) The graphs report the mean±SEM of the number of filopodia per 100µm of circumference, their length, width, lifespan and solidity, respectively. **F**) The graph reports the mean±SEM of the percentage of the frequency distribution of the lifespan of filopodia. A y-axis truncated version is shown to highlight the small fractions of filopodia with lifespans exceeding 90min. A full version of the graph is shown in Fig. S4A. The multiple Mann-Whitney tests across time points show significant differences between WT and EFA6B KO spheroids at 360min with *p* value of 0.0016, as well as between WT and EFA6B KO+GM6001 spheroids with *p* value of 0.0067. No significant difference between untreated and GM6001-treated EFA6B KO spheroids. In B, C, E, F and G: 1711, 2481 and 2376 filopodia were analyzed from 10 WT, EFA6B KO and EFA6B KO+GM6001 spheroids from three independent experiments, respectively. *P* values calculated using an unpaired t test, \**p*<0.05; \*\**p*<0.01; \*\*\**p*<0.005 and n.s.: not significant. **H**) Fixed phase contrast and fluorescent (nucleus) images from one representative movie (see four additional examples (Fig. S5C-F as well as the original movies S2-S6) of a MCF10 EFA6B KO invasive spheroid embedded in collagen I gel (2mg/ml) showing a stable nascent filopodia, the same filopodia just before and after nuclear translocation. The width in µm of the filopodia are indicated. Scale bar 25µm. **I**) The graph reports the mean±SEM of the width of the filopodia at each step for five spheroids from two independent experiments. *P* values calculated using an unpaired t test, \**p*<0.05; \*\**p*<0.01.

## Discussion

The precise mechanisms of BM infiltration by tumor cells remain poorly understood due to the lack of reliable *in vitro* models and the challenges associated with *in vivo* and *ex vivo* investigations. While Matrigel and synthetic BM substitutes are useful for assessing specific BM properties, such as stiffness and elasticity, and for evaluating certain aspects of cell invasiveness, they do not fully recapitulate the complex composition and organization of physiological BM. Studies utilizing animal models of tumor cell invasion remain challenging due to the labor-intensive, time-consuming, and costly nature of imaging and genetic manipulation. Here, we have characterized cell spheroid models that autonomously synthesize and assemble a BM exhibiting all the key attributes of a normal BM. This approach enables high spatiotemporal resolution analysis of BM assembly in 3D culture systems and makes possible genetic manipulation, as well as pharmacological, biochemical, or molecular perturbations.

The BMs assembled by our cell models, MCF10 and HMLE, are unlikely to perfectly replicate the BM of the human mammary gland, which itself varies depending on breast development and its localization around lobules or ducts. However, they contain all the essential core components, including laminins, collagen IV, cross-linkers (nidogen, perlecan), and hemidesmosomes. Also, the EFA6B mutation found in patients with aggressive breast tumors facilitates invasion *in vitro* and in an *in vivo* model of DCIS-to IDC transition (17, 18). Therefore, they serve as suitable models to study the BM architecture, functions and infiltration. Another benefit of our invasive models is that the EFA6B mutated organoids can be compared with their corresponding wild-type controls, which is not possible when using classical invasive human cell lines. Our experimental approach can be adapted to other cell types as well as co-culture systems to further increase complexity. It enabled us to investigate in an accessible and controlled *in vitro* set up the BM architectural organization and the mechanisms of its infiltration by tumor cell models.

We first studied the overall architecture of the BM. By studying both cell spheroid models, we observed that all major components of the BM were not only distributed along a common lattice but also exhibited an almost continuous organization. Although laminin γ2 staining appeared more as small puncta, it was detectable throughout the BM lattice, indicating that small hemidesmosomes may form in a dense and widespread manner. Similarly, perlecan staining appeared nearly continuous, suggesting that cross-linking may tightly be maintained along the entire BM lattice rather than as spread sites of connections. Additionally, we observed that the common lattice contained pores with a surface area in the range of 0.6µm², raising questions about the molecular arrangement that accommodates these pores. Given the estimated sizes of collagen IV molecules and that of the N-terminal domains of the laminin chains, they are predicted to form lattices with surface areas of approximately 0.5µm² and 0.005µm², respectively (31, 32). Thus, we propose that the collagen IV meshwork outlines the pores. Further, since both meshworks appear superimposed, our observations suggest that the collagen IV meshwork determines the overall lattice structure and that a regulatory mechanism must exist to confine the laminin network to align along the collagen IV framework.

After characterizing the BM assembled by our 3D mammary spheroid models and detailing its architecture, we next investigated its infiltration. In a previous study, we reported that gene set enrichment analysis (GSEA) signatures associated with the extracellular matrix (ECM), including the basement membrane (BM) signature, were among the most affected when comparing non-invasive ductal carcinoma in situ (DCIS) to invasive ductal carcinoma (IDC) in human breast tumors (18). Given that tumor cells have been shown to contribute components to the BM (14, 34), we hypothesized that invasive cells, with an altered repertoire of BM components might promote the organization of a BM more permissive to infiltration. To test this hypothesis, we took advantage of our EFA6B KO cell invasion model, which displays the same alterations in the ECM and BM gene signatures as those observed during the DCIS to IDC transition in tumor patients. Thus, we examined BM assembly in the MCF10 and HMLE EFA6B KO cell models. In both cases, we observed a reduction in BM component secretion and a decreased assembly on the surface of the spheroids. The most striking observation was the substantial increase in pore size. Notably, many of the pores exceeded 2µm^2^, a size sufficient to permit cell escape (35–37). The average pore area doubled, consistent with the loss of a node within the collagen IV polygonal organization, leading to the fusion of pores.

The pores in the BM are not empty spaces. The 3D reconstruction of serial section images obtained via confocal microscopy revealed that all pores were filled by plasma membrane extensions. The extension of filopodia forming large pores through the BM challenges the concept of its nanoporous nature (as observed by SEM or inferred from small-molecule diffusion studies (9)). We propose that the nanoporous organization observed by SEM is the result of a staggering arrangement of multiple layers of flexible collagen IV networks (38). However, the extension of filopodia may locally realign these overlapping collagen IV lattices, creating a pore with the molecular dimension corresponding to that of a collagen IV parallelepiped of about 0.5µm^2^.

In invasive EFA6B KO cell spheroids, the number and length of filopodia traversing the BM were markedly increased. Concurrently, the BM exhibited larger pores, suggesting that the filopodia might contribute to pore enlargement. To test this hypothesis, we induced filopodia extension in cells at the periphery of the spheroids and measured the impact on BM pore size. Previous studies have reported that the myosin II inhibitor blebbistatin stimulates the formation of long filopodia in MCF10 cells (13). Upon exposure to blebbistatin, we observed a significant induction of filopodia formation, which was accompanied by an increase in BM pore size. Further, comparative analysis of the MCF10 cell line and its invasive derivative, DCIS.com, has revealed that DCIS.com cells exhibit a higher density and length of filopodia. We now report that the increased filopodia activity is associated with larger pores in the BM of DCIS.com spheroids compared to MCF10WT spheroids. Collectively, these findings indicate that filopodia produced by invasive cells caused enlargement of BM pores.

We showed that the BM surrounding spheroids of invasive cells contained larger pores, which were generated by the extension of longer filopodia. Next, we aimed to determine whether the largest pores, exceeding 2µm in size, served as exit points for cell infiltrating the BM. Using videomicroscopy, we observed that spheroids actively projected and retracted numerous thin filopodia; however, cell escape occurred exclusively at sites where a large filopodia had previously extended. In fact, cell exit was consistently preceded by the formation of a thick filopodium, suggesting that it created a sufficiently large opening in the BM to facilitate cell escape. Further, these large filopodia could be distinguished from the numerous thinner ones by their sensitivity to the protease inhibitor GM6001, suggesting that protease activity is required to facilitate the breakdown of the BM meshwork and the formation of a large pore. A similar pattern of filopodial-mediated invasion through reconstituted BM and alginate-based hydrogels was observed (39). To the best of our knowledge, we present the first evidence that long protrusive filopodia, known to be extended by cancer cells (33), perforate a naturally assembled BM to allow for the escape of invasive cells, in a mechanism that is reminiscent of the *C. elegans* anchor cell (12).

The cellular process by which metastatic cells traverse the BM remains poorly understood. Several mechanisms have been proposed: enzymatic degradation of the BM by proteases (40), mechanical forces exerted on the BM by tumor cells or cancer-associated fibroblasts (CAFs) (11, 41), and alterations in the molecular composition of the BM that modify its biophysical properties (14). It is unlikely that a single mechanism alone is sufficient to overcome this barrier. We propose a model in which tumor cells first modify the composition of the BM, rendering it more susceptible to a combination of forces generated by tumor growth and filopodia-driven remodelling of its molecular architecture. These structural alterations would facilitate protease activity, leading to the formation of larger openings in the BM. Further research is required to characterize the alterations in the BM that increase its permissiveness to invasion, identify the molecular players involved in the mechanical forces exerted on the BM, and determine the specific proteases that create structural vulnerabilities. These findings would need to be validated using *in vivo* models that allow real-time tracking of tumor cells and BM dynamics during infiltration.

## Materials and Methods

### Cells and reagents

MCF10A cells were obtained from ATCC (LGC Standards, France) and grown in DMEM/F-12 (1:1), horse serum 5%, non-essential amino acids 1%, insulin 10μg/ml, hydrocortisone 1μg/ml, EGF 10ng/ml, cholera toxin 100ng/ml and penicillin (100u/ml)-streptomycin (100μg/ml). HMLE cells were obtained from Dr. R.A. Weinberg (Whitehead Institute for Biomedical Research, Cambridge, MA, USA) (42) and grown in DMEM/F-12 (1:1), fetal calf serum 10%, insulin 10μg/ml, hydrocortisone 0.5μg/ml and penicillin (100u/ml)-streptomycin (100 μg/ml). DCIS.com cells were obtained from Dr. P. Chavrier (Institut Curie, Paris, France) and grown in DMEM/F-12 (1:1), L-glutamine 2mM, horse serum 5%. All culture reagents were from Invitrogen (Fisher Scientific, France) except for the fetal calf serum (Dutscher, France). The PSD4/EFA6B KO MCF10 and HMLE were previously described (18). All secondary antibodies and fluorescent probes were from Molecular Probes (Invitrogen). For the list of primary antibodies refer to Table S1. Unless otherwise indicated, all other reagents were from (Sigma-Aldrich, France).

### 3D culture

3D culture was performed using Matrigel (5mg/ml) or rat tail Collagen I (Corning^®^, Fisher Scientific, France). The collagen I solution was neutralized using 1N NaOH and diluted in PBS to a final concentration of 2mg/ml. 5µl of 0,5x10^6^ cells was mixed in a 25µl drop of Matrigel deposited on a glass coverslip in 24-well plate, or in 150µl of Collagen I placed in a well of an 8-well Nunc™ Labtek™ dish. For SEM and AFM, spheroids prepared in Matrigel were extracted as previously described (43). Briefly, 2.5x10^5^ cells mixed in 300µl of ice-cold Matrigel was deposited in a 24-well plate and placed at 37°C. After 30min, 500µl of Matrigel was added on top and the spheroids let grown for 4 days. For the extraction, after 2 quick washes in ice-cold PBS, the sample was incubated 30min at 4°C in 0.5ml PBS-EDTA (5mM) under agitation. The liquefied gel was then transferred into a 1.5ml Eppendorf tube, centrifuged at 0.3g for 2.5min, the supernatant discarded and the pelleted spheroids washed once in ice-cold complete medium and centrifuged at 0.3g for 2.5min. The supernatant was discarded, the spheroids gently resuspended in ice-cold complete medium and deposited on a glass coverslip or dish for SEM and AFM analyses, respectively.

### Immunoblot

To prepare whole cell lysates, cells grown on plastic dishes washed with PBS were lysed with an SDS lysis buffer (0.5% SDS, 150mM NaCl, 5mM EDTA, 20mM Triethanolamine–HCl pH 8.1, 1mM PMSF). The lysate was heated at 95°C for 10 min and then thoroughly vortexed for 15 min. After centrifugation at 16,000 *g* for 20 min at room temperature, the supernatant was transferred into a new tube containing 5× Laemmli buffer and further boiled 5 min at 80. To recover proteins from the culture supernatants, two 10-cm Petri dishes were plated with 4x10^6^ cells. The next day, the cells were extensively washed with PBS and incubated 48hr in 9ml serum-free culture medium. The supernatants were cleared by high-speed centrifugation, and then concentrated using Amicon® Ultra-15 (100kDa MW) units (Sigma-Aldrich) down to 250µl to which 50µl of 6X Laemmli buffer was added before 3min boiling at 80°C. Equal amounts of whole cell lysates and concentrated supernatants were loaded into SDS-PAGE and transferred onto a nitrocellulose membrane. Membrane blocking and secondary antibodies dilutions were done in PBS 5% non-fat dry milk, primary antibodies were diluted in PBS 3% BSA. The proteins were revealed by chemiluminescence (ECL™, Amersham France) using secondary antibodies directly coupled to HRP. The membranes were analyzed with the luminescent image analyzer Fusion (VILBER, France) and band intensity quantified using the image analyzer software AIDA (Elysia-raytest, Germany). For quantitative analysis, results were normalized to the loading controls and to the control sample arbitrarily set at 1.

### Mass Spectrometry analysis

10µg of proteins was loaded on precast Tris-Glycine gels (Bio-Rad, France), and stained with Coomassie blue (Thermo Fisher Scientific, France). Protein spots were excised from the gel, distained by adding H_2_O/acetonitrile (1/1), rinsed (15 min) in acetonitrile and dried under vacuum. Each excised spot was reduced in dithiothreitol, alkylated with iodoacetamide and washed successively in H_2_O/acetonitrile (1/1), and acetonitrile. Next, gel pieces were digested in NH_4_HCO_3_ buffer containing 10ng/mL of trypsin (modified porcine trypsin sequence grade, Promega) for one hour at 4°C, and OVN in NH_4_HCO_3_ buffer without trypsin at 37°C. Tryptic peptides were isolated by extraction with 60µL of 1% acid formic and 60µL acetonitrile. Peptide extracts were pooled, concentrated under vacuum, solubilized in 15µL of aqueous 0.1% formic acid and separated on a nanoHPLC (ultimate 3000, Thermo Fisher Scientific). 5µL of peptidic solution was injected and concentrated on a µ-Precolumn Cartridge Acclaim PepMap 100 C18 (i.d. 5mM, 5µm, 100 Å, Thermo Fisher Scientific). Peptides separation was performed on a 75µm i.d. x 500mM (3µm, 100 Å) Acclaim PepMap 100 C18 column (Thermo Fisher Scientific) at a flow rate of 200nL/min. Solvent systems were: (A) 100% water, 0.1% formic acid, (B) 100% acetonitrile, 0.08% formic acid. The nanoHPLC was coupled via a nanoelectrospray ionization source to a Hybrid QuadrupoleOrbitrap High Resolution Mass Spectrometer (Thermo Fisher Scientific). MS spectra were acquired at a resolution of 70 000 (at 200m/z) with a scan range of 150-1800m/z, an AGC target value of 5e5 and a maximum injection time of 50ms. 10 most intense precursor ions were selected and isolated with a window of 2 m/z and fragmented by Higher energy C-Trap Dissociation with normalized collision energy NCE of 27. MS/MS spectra were acquired in the ion trap at a resolution of 17 500 (at 200 m/z) with an AGC target value of 2e5 and a maximum injection time of 100ms.

### Mass spectrometry data processing and statistical analysis

MS data were subjected to LFQ analysis using the MaxQuant platform (v2.0.3.1) (http://www.maxquant.org/). Database search of the MS/MS data was performed in MaxQuant using the Andromeda search engine against Human reviewed UniProtKB database (july 2020) and MaxQuant contaminants database. Digestion mode was set to Trypsin/ P specificity, with a fixed carbamidomethyl modification of cysteine, and variable modifications of protein N-terminal acetylation and methionine oxidation. Mass deviation was set to 20 ppm for the first and 6 ppm for the main search and the maximum number of missed cleavages was set to 2. Peptide and site false discovery rate (FDR) were set to 0.01. Proteins identification was performed using a minimum of 1 unique+razor peptide. Quantification was achieved using the LFQ (Label-Free Quantification) algorithm. Unique+razor peptide was used for LFQ quantification and the “label min. ratio count” was fixed at 1. The match between runs option was enabled, allowing a time window of 0.7 min to search for already identified peptides in all obtained chromatograms.

MaxQuant-processed data were loaded in R (v4.3.2) as text-files. After filtering out decoy and contaminant matches, peptides without quantification in more than 1 sample upon the 3 replicates for both conditions were discarded. Using the Limma R package (v3.56.2), a linear model was fitted to log2-transformed protein intensities, and contrasts were applied to compare MCF10 EFA6B KO and WT conditions. Empirical Bayes smoothing was then used to obtain stable differential expression estimates. The DEqMS package (v1.18.0) extends the Limma framework by adjusting variance estimates based on the number of peptides or peptide-spectrum matches (PSMs) per protein, ensuring accurate differential expression analysis. The minimum peptide counts for each protein, with a pseudocount of 1 added to avoid zero values, was incorporated into the model. The spectraCounteBayes function was applied to correct bias in variance estimates, resulting in a differential protein abundance table. This table includes, for each protein, the log fold change and a spectra count-adjusted posterior p-value, corrected for multiple comparisons using the Benjamini-Hochberg method. The UniProt ID Mapping tool was used to retrieve Ensembl gene symbols from UniProt protein IDs. Gene symbols for significantly upregulated (sca.adj.pval < 0.05 and logFC > 0.25) and downregulated (sca.adj.pval < 0.05 and logFC < -0.25) proteins were then grouped into separate gene sets. Gene Set Enrichment Analysis (GSEA) was performed using EnrichR on the Gene Ontology collection to identify pathways significantly modulated by the EFA6B KO.

### Joint analysis of publicly available scRNA-seq datasets of human mammary cell lines and physiological tissue

Raw gene counts matrices from MCF10 and HMLE cell lines (19, 20) were filtered to retain only untreated cells from both studies (n = 1337 cells and n = 1948). To manage computational load, a random 10% subset of cells from the full Human Breast Cell Atlas dataset was selected (n = 51209 cells). Cell type annotations and raw counts from these datasets were integrated with those from the cell lines, resulting in a unified annotated data matrix processed using the Seurat R package (v4.3.0.1). For gene expression representation, raw counts were normalized and log2-transformed using the NormalizeData function.

### Immunofluorescence

After three washes in PBS, the spheroids embedded in Matrigel or Collagen I gels were fixed in 4% paraformaldehyde for 30 min, washed 3 times and then incubated in PBS containing 2% fish skin gelatin. Then the cells were incubated with primary antibodies over-night and counterstained with appropriate fluorescent secondary antibodies, DAPI and phalloidin as indicated in the figure legends. Images acquisition was performed with a confocal microscope TCS SP8 with a HCX PL APO 63X/1.4 objective (Leica Microsystems). For Maximum Intensity Projection (MIP) and 3D reconstruction samples were scanned every 200nm with an optical section of 896nm for a wavelength of 580nm.

### Pore detection and area quantification

To perform the pores detection, we considered 2D slices due to the difference between XY and Z resolution. After a comparison study between a marked point process based on algorithm (44) and the convolutional networks Stardist (https://github.com/stardist/stardist) and Cellpose (https://www.cellpose.org/), we selected Cellpose as the most promising model. In order to improve the model performances, we fine-tuned the parameters by annotated 12 slices using the software LabKit. We considered a pre-processing step consisting of a gaussian filtering. The model was trained and the parameters fixed following a cross-validation scheme where 25% of the samples were used for training (i.e. 4 slices) and 75% for the validation. To estimate the performances, we split the test set into two subsets, “easy” and “difficult”, depending on the noise. The results and an example are shown in Fig. S5A,B. After detecting the pores in 2D, we reconstructed the 3D objects by connecting pores in adjacent slices. Note that we improved the detection precision as some pores can be misdetected in one slice but detected in the adjacent slice. To compute the size of each pore we projected the detected object onto the spheroid tangent plane. To estimate this plane, we considered the alpha shape of the whole spheroid obtained by a threshold (45).

### Quantification of filopodia from fixed spheroids embedded in Matrigel

Filopodia of fixed spheroids were quantified using the Single Image Filoquant plugin (33) implemented on Fiji according to the instructions of the authors. For an unbiased quantification the entire periphery of the spheroids was analysed and the same settings used for all images. 10 spheroids from 3 independent experiments were analyzed in a randomized manner by a blind third party.

### Scanning electron microscopy

Spheroids were prepared in Matrigel 5mg/ml, extracted at day 4 as described above and deposited on a glass coverslip overnight. After fixation by immersion in 2.5% glutaraldehyde in 0.1M cacodylate buffer pH7.4, the samples were rinsed in buffer, dehydrated in a graded ethanol series, and finally immersed in hexamethyldisilazane (Carl Roth, Karlsruhe, Germany), and dried at room temperature. Samples were then mounted on aluminium stubs and sputter-coated with a 3-nm gold-palladium coating (Cressington 308EM, UK) prior to analysis with a Field Emission Scanning Electron Microscope (FESEMJEOL 6700F, Japan).

### Transmission electron microscopy

Spheroids were grown in Matrigel 5mg/ml for 4 days, then immerged in fixative, 2.5% glutaraldehyde in 0.1 M cacodylate buffer pH7.4 and stored overnight at 4°C. Samples were rinsed in the same buffer, post-fixed for 1h30 in 1% osmium tetroxide and 1% potassium ferrocyanide in 0.1M cacodylate buffer pH7.4 to enhance the staining of membranes. Samples were then rinsed in distilled water (overnight at 4°C), dehydrated in acetone and lastly embedded in epoxy resin. Air bubbles were removed by several vacuum cycles. Classically contrasted ultrathin sections (70 nm) were analysed under a JEOL 1400 transmission electron microscope equipped with a Morada Olympus CCD camera.

### Atomic Force microscopy

Spheroids grown in Matrigel were extracted as described above and plated overnight in a 50mm glass bottom dish (WillCo Wells, Netherlands) in complete medium. The next day the immobilized spheroids were washed and placed in Leibovitz’s medium supplemented with 2% horse serum. The sample quantitative mechanical properties were obtained using a Bioscope Catalyst operating in Point and Shoot mode (Bruker Nano Surfaces, Santa Barbara, CA, USA), equipped with a Nanoscope V controller and coupled with an inverted optical microscope (Leica DMI6000B, Leica Microsystems Ltd., UK). The force-distance curves were captured using a Borosilicate Glass spherical tip (5μm of diameter) mounted on a cantilever with a nominal spring constant of 0.06N/m (Novascan Technologies, Ames, IA USA) with a trigger threshold of 1nN and a ramp velocity of 2μm/s. Before performing the experiments, the cantilever was left for at least 20 minutes in the medium environment to allow its thermal stabilization prior to calibrate the optical lever sensitivity against a clean Willco dish and calculate the actual cantilever spring constant by using the thermal tune method. The apparent Young’s modulus was calculated using the NanoScope Analysis 1.80 software (Bruker Nano Surfaces, Santa Barbara, CA, USA) applying to the force curves, after the baseline correction, the Hertz spherical indentation model using a Poisson’s ratio of 0.5. To avoid any tilt effect due to the base correction step, only the force curves having their maximum value at 1nN were taken into account for performing the fit, which had the force fit boundaries between 0% and 25% (corresponding to a maximal force of 250pN). Among the processed force-distance curves, only the one having an apparent Young’s modulus values corresponding to a fit with R2> 0.80 were taken into account. For all samples, in each independent experiments, from 3 to a maximum of 19 force curves/spheroid, were used for calculating the corresponding Young’s modulus (YM) values (n). The superplot (GraphPad Prism 10.4.1) is expressed as a scattered plot, in which each dot (n) corresponds to the YM mean value/spheroid having a different color for each independent experiment (N). Every triangle corresponds to the YM mean ± SD, calculated from all the YM mean value/spheroid (n) corresponding to each independent experiment (N). A paired t-test, applied to the YM mean values obtained for each N was used.

### Time-lapse imaging

To form spheroids, 50x10^3^ cells were incubated in complete medium for 48h in poly-HEMA coated 35mm dishes. Size homogenous spheroids isolated by differential low-speed centrifugation were resuspended in 300μl of fibrillary collagen I (2.5mg/ml) and plated in 24-well plates. 48h later images were acquired using a Cytation^TM^ 5 imaging multi-mode reader (BioTek Instruments) in a controlled environment. Time-lapse images were captured every 30min and up to 72h with a 20X/0.45 phase contrast objective using the Gen5 v3.13 software (Biotek Instruments). For nuclei live staining, Hoechst 33342 was added at 5nM in complete medium 16h before imaging.

### Detection and quantitative analyses of filopodia in spheroids videomicroscopy

We address the filopodia detection in 2D by first normalizing each slice between 0 and 255. We observed that the background of the image was subject to illumination variations that correspond to low frequencies. To estimate this low frequency component, we convolved the image with a gaussian filter of high variance *G*. We then considered *W = G-I* and *B = I-G*, filtered using a median filter to reduce the noise, for the detection of the spheroid and the filopodia, respectively (**Fig. S5C**).

The first consisted in detecting the spheroid. Visually this object was not defined by its intensity properties but rather by some specific texture. To detect this texture with some localization accuracy, we computed the conditional variance in local windows (46). A threshold is applied to obtained the texture object. We then selected the biggest connected component (corresponding to the spheroid and adjacent filopodia), filled the holes and applied an opening to remove the filopodia. Finally, we fitted an ellipse *E* to the detected object in order to approximate the spheroid (**Fig. S5D**).

To detect the filopodia we first applied a diffusion filter to both *W* and *B* to reduce the residual noise. We then thresholded both images and masked the detected ellipse MW=(W>T)*(1-E), MB=(B>T)*(1-E). Labelling the connected components of both MW and MB identified filopodia candidates. A candidate was validated if: its size was between minimum and maximum thresholds, its orientation was close to the spheroid normal at the spheroid closest point, its distance to the spheroid was below a threshold. After this validation process we still may obtain multiple detections (black and white) due to the diffraction halo. We therefore used the following rules: i) if a white filipodium was close to two black filipodia sharing a similar orientation, we removed the two black filipodia; ii) if a black filipodium was close to two white filipodia sharing a similar orientation we removed the two white filipodia; iii) if two filipodia, respectively black and white were close and shared a similar orientation we kept the white one (**Fig. S5E**). Finally, we reconstructed the time stack and labelled in 3D the connected components. As output, we extracted for each detected filipodium the following parameters: i) its localization w.r.t the spheroid (angle); ii) time of arrival and time of departure; iii) maximum length, iv) width (see below) and v) solidity (ratio of the pore size and its convex envelope size).

Due to halos created around the filopodia by the diffraction of light, our custom software was unable to accurately determine the width. Therefore, measurements were performed manually using Gen5 v3.13 software by a blind third party. The lifespan was estimated by adding up the number of 30-minute increments during which filopodia were detected.

### Statistics

All experiments were performed at least three times independently. The number of cells, spheroids, pores or filopodia analyzed are indicated in the legends. Error bars represent ± standard error of the mean (SEM). Statistical significance was determined using a Student t-test and one-way ANOVA, in which *p* values <0.05 were considered statistically significant. All statistical analyses were performed with GraphPad Prism software version 10.4.1.

## Acknowledgments

This work has been supported by the French government through the UCA^JEDI^ Investments in the Future project managed by the National Research Agency (ANR) with the reference number ANR-15-IDEX-01. We warmly thank Alexandre Helpiquet and Kelly Cappelli for preliminary experimental work and Dr. Karen Singer for critical reading of the manuscript.

---

**Fig. S1.**
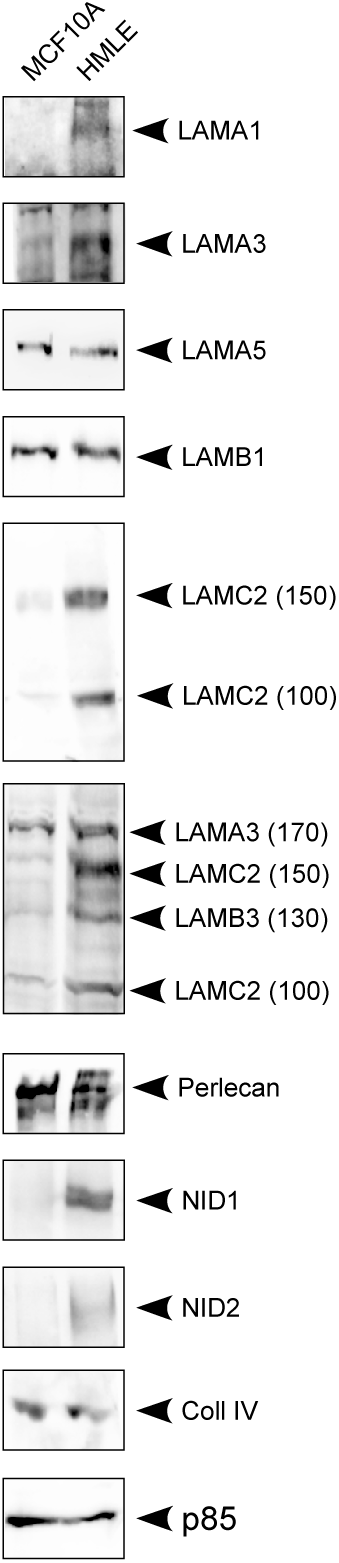
MCF10 and HMLE cells were solubilized and the expression of the indicated proteins analyzed by immunoblot. p85 served as a loading control.

**Fig. S2.**
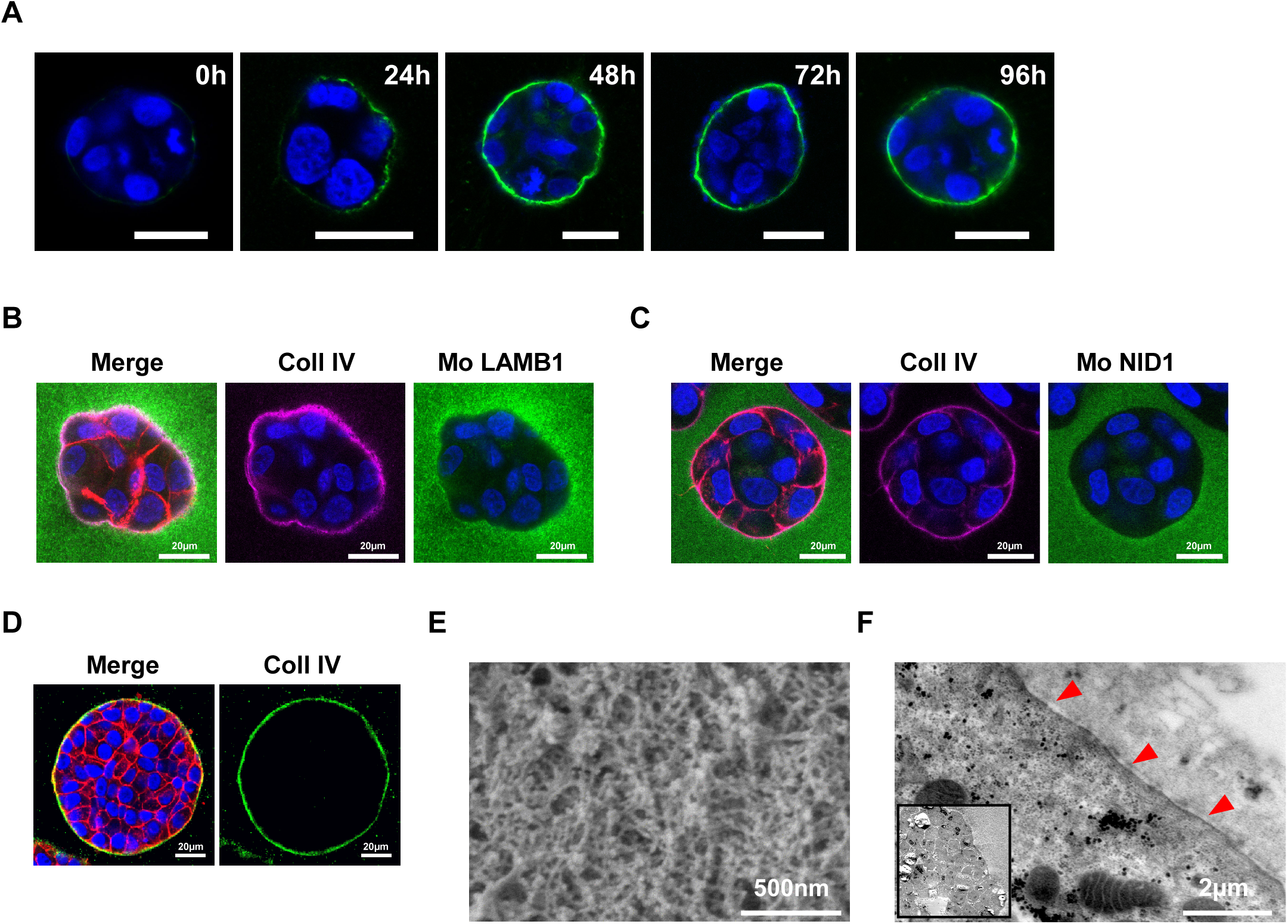
**A**) Representative confocal immunofluorescence images of the time course of the BM assembly at the surface of a MCF10 spheroid followed overtime by collagen IV staining (green). DAPI in blue. Scale bar 20µm. **B**) and **C**) Representative confocal immunofluorescence images of MCF10 spheroids stained for collagen IV (magenta), nuclei (blue) and murine LAMB1 or murine Nidogen 1 (green). **D**) Representative confocal immunofluorescence images of a MCF10 spheroid grown 120h in collagen I gel and stained for collagen IV (green), F-actin (red), nuclei (blue). **E**) SEM image of a MCF10 spheroid. **F**) TEM image of the surface of an HMLE spheroid (inset: whole spheroid). Red arrows point to the thin dark line formed by the BM lining the spheroid.

**Fig. S3.**
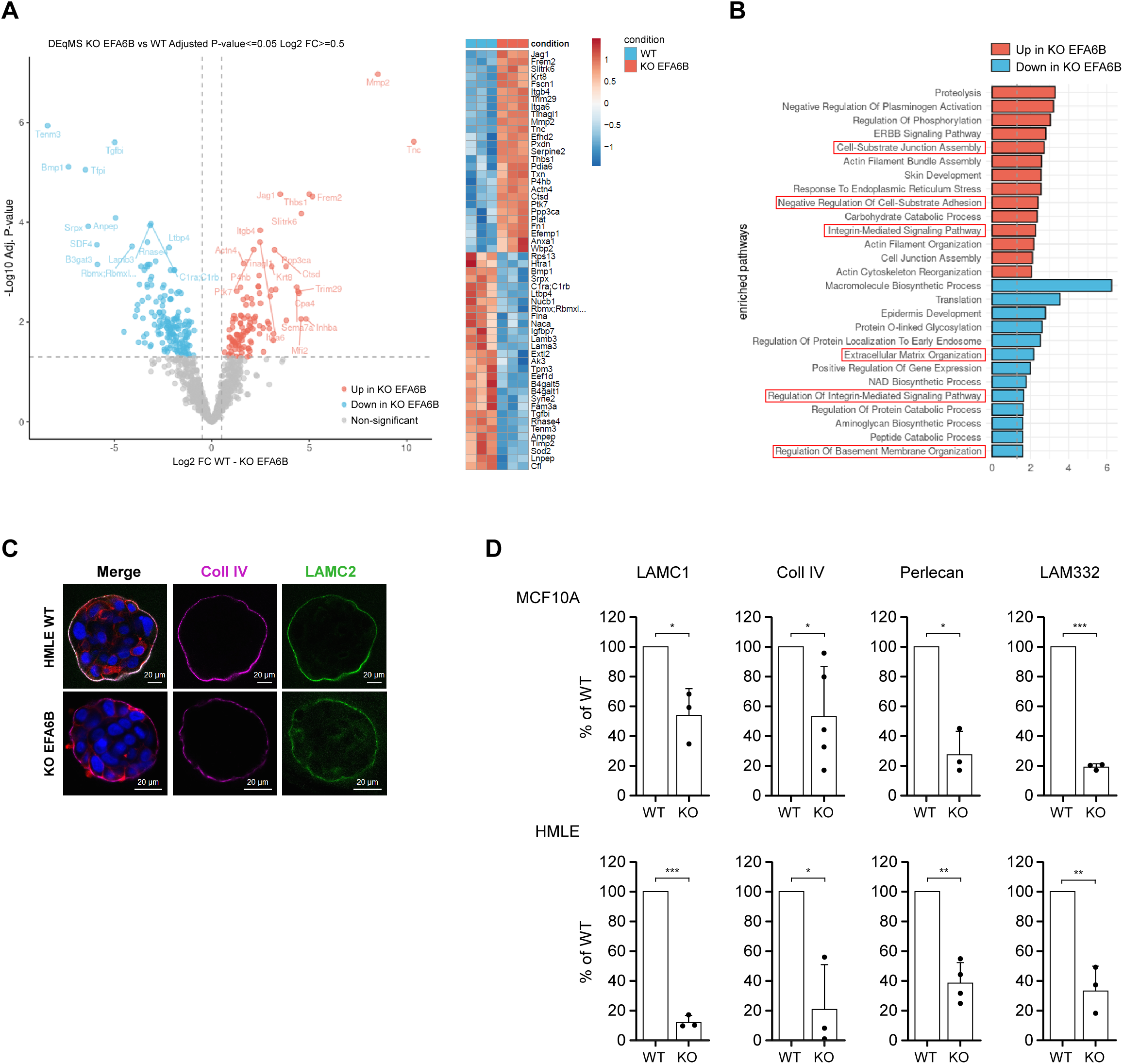
**A**) Volcano plot of the differential protein abundance of the BM components in MCF10WT and EFA6B KO cells (left panel). Heatmap of z-scored LFQ intensities for the most affected proteins (right panel). **B**) Gene set enrichment analysis (GSEA) of the significantly up and down-regulated proteins by EFA6B KO in MCF10 cells. Circled in red GO related to the ECM, the BM and the integrin-mediated interaction are affected. **C**) Representative confocal immunofluorescence images of HMLE WT (top) and EFA6B KO (bottom) spheroids grown 96h in Matrigel and co-stained for Coll IV (magenta), LAMC2 (green), F-actin (red), nuclei (blue). **D**) Quantitative analysis of the immunoblots shown in Figure 3B. The results were normalized to the WT cells. N=3-5 as indicated on the graphs. P values calculated using a paired t test with Prism software. *p<0.05; **p<0.01; ***p<0.005.

**Fig. S4.**
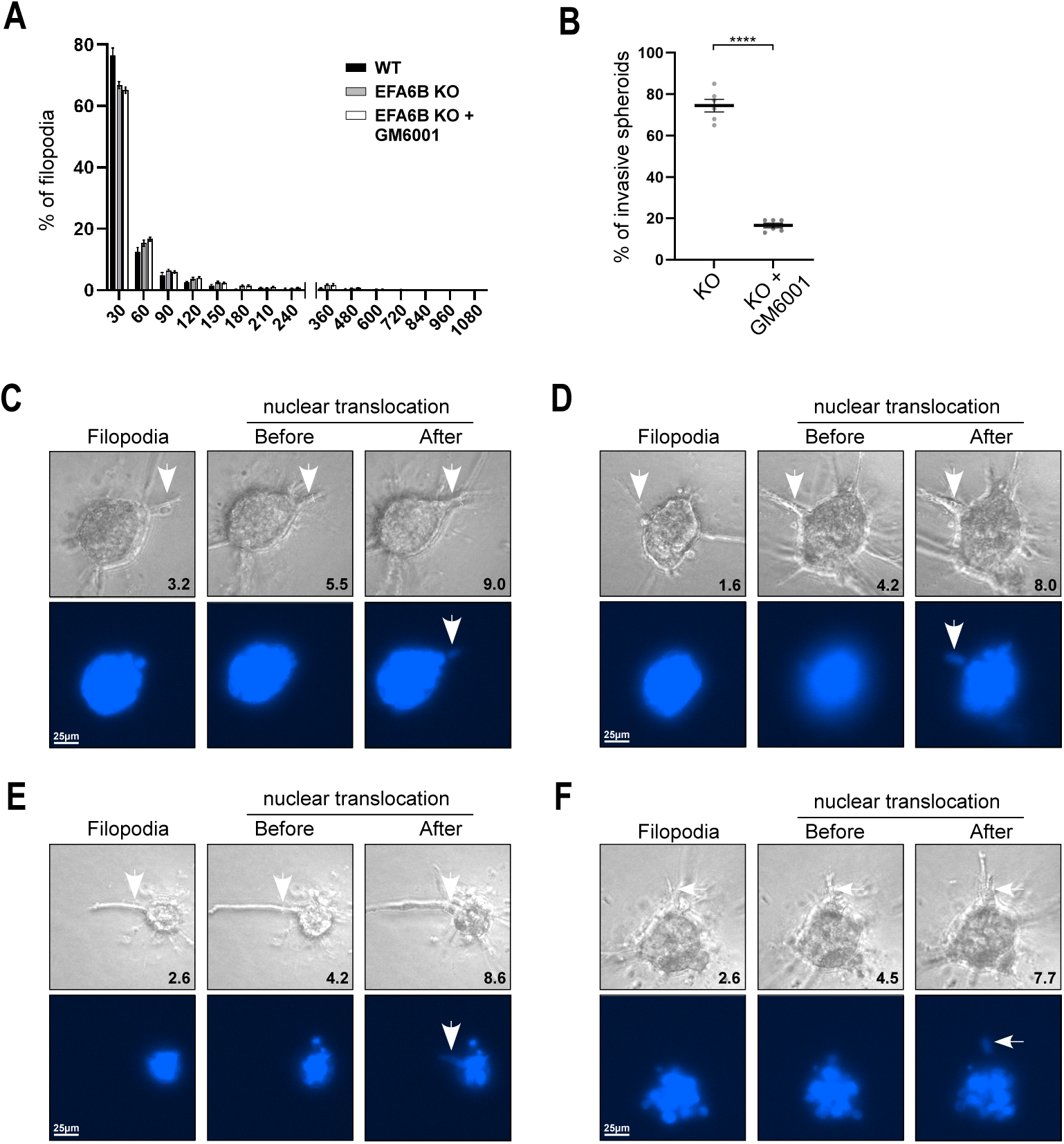
**A**) The graph reports the mean±SEM of the percentage of the frequency distribution of the lifespan of filopodia. **B**) Quantification of the percentage of cell aggregates (n =100) with invasive protrusions of EFA6B KO cell spheroids treated or not with GM6001(25µM) grown in collagen for 3 days. N = 3, average ± SEM. P value calculated using a paired t test, ****p<0.0001. **C-F**) Fixed phase contrast and fluorescent (nucleus) images from four movies (Movies S3-S6) of MCF10 EFA6B KO invasive spheroids embedded in collagen I gel (2mg/ml) showing a stable nascent filopodia, the same filopodia just before and after nuclear translocation. The width in µm of the filopodia are indicated. Scale bar 25µm.

**Fig. S5.**
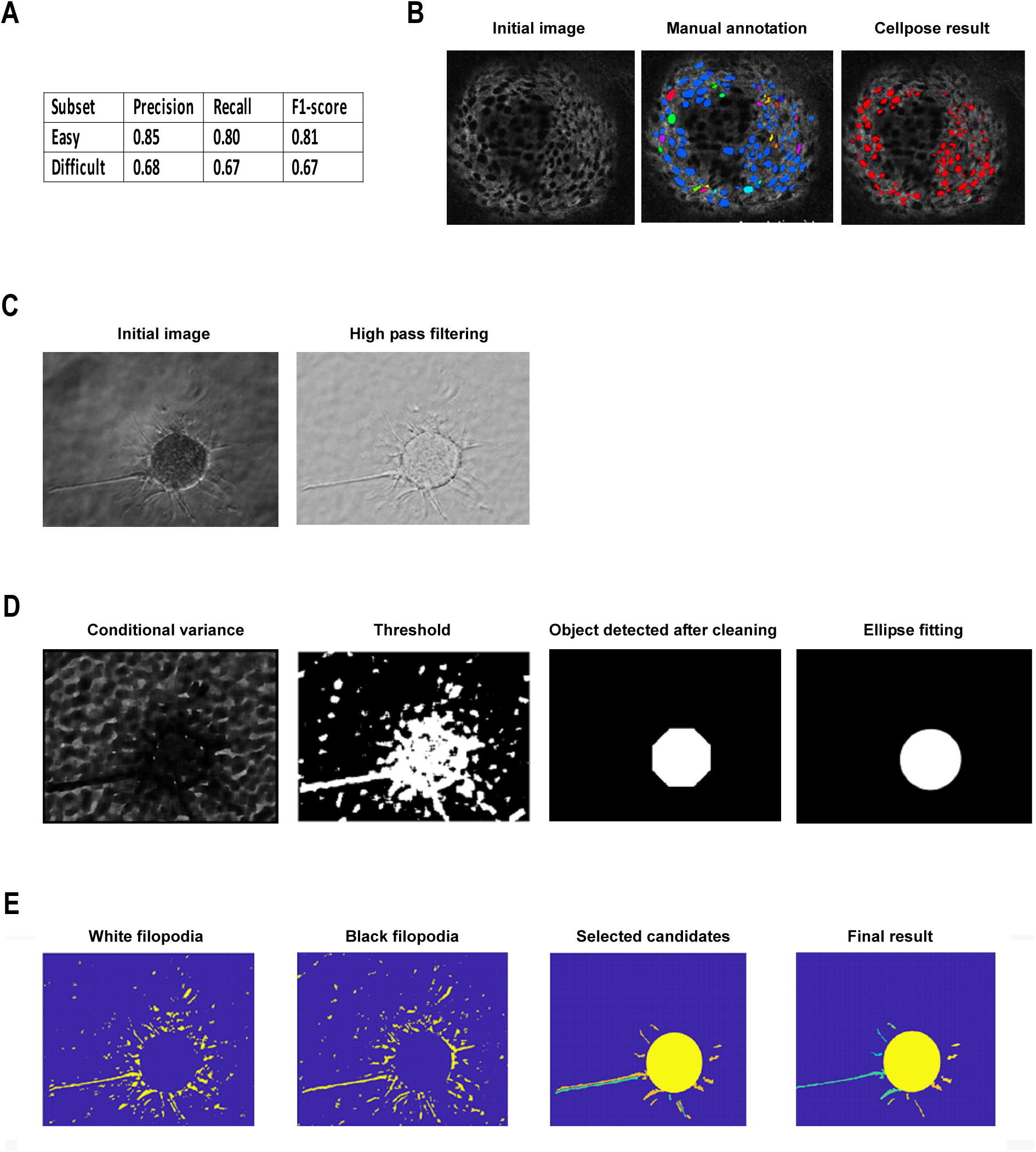
**A**) Cellpose performance on 2D slices **B**) Example of the manual and Cellpose detection of pores. **C**,**D**,**E**) Images illustrating the sequential steps of the custom-made software for sharping the image (C), detecting the spheroid (D) then its filopodia (E, white in orange and black in turquoise).

**Table S1:** List of primary antibodies used in this study.

| Antigen | Antibody source and reference | Application (dilution from stock or concentration) |
| --- | --- | --- |
| LAMA1 | Santa Cruz, clone G12 | Immunoblot 1/500 |
| LAMA3 | mAb BM165<sup>1</sup> | Immunoblot 1/500 |
| LAMA5 | Bio-Rad, clone 4C7 | Immunofluorescence 1/50 |
| LAMA5 | Invitrogen, clone CL3118 | Immunoblot 1/500 |
| LAMB1 | Santa Cruz, clone D9 | Immunoblot 1/500 |
| LAMB1 (Mouse) | R&D, clone AL-3 | Immunofluorescence 1/50 |
| LAMC1 | Novus Bio, clone A5 | Immunofluorescence 1/50 |
| LAMC1 | Santa Cruz, clone B4 | Immunoblot 1/500 |
| LAMC2 | Millipore, clone D4B5 | Immunofluorescence 1/50 |
| LAMC2 | Santa Cruz, clone B2 | Immunoblot 1/500 |
| LAM332 | pAb 4101<sup>1</sup> | Immunoblot |
| Coll IV | Millipore, AB769 | Immunofluorescence 1/50 |
| Coll IV (COL4A1) | Cell Signaling, 50273 | Immunoblot 1/500 |
| Perlecan | Santa Cruz, clone E6 | Immunofluorescence 1/50 |
| Perlecan | Santa Cruz, clone A7L6 | Immunoblot 1/500 |
| NID1 | R&D, clone 302117 | Immunofluorescence 1/50<br>Immunoblot 1/500 |
| NID1 (Mouse) | Millipore, clone ELM1 | Immunofluorescence 1/50 |
| NID2 | Santa Cruz, clone F2 | Immunoblot 1/500 |
| Actin | Sigma, A4700 | Immunoblot 1/500 |
| Actinin | Sigma, A5044 | Immunoblot 1/500 |
| GST | GE Healthcare, 27-4577-01 | Immunoblot 1/1000 |
| Hsp60 | Sigma, SAB4501464 | Immunoblot 1/200 |
| p85 | Millipore, ABS1856 | Immunoblot 1/200 |
| GAPDH | Sigma, G9545 | Immunoblot 1/500 |

1. Bachy, S., Letourneur, F. & Rousselle, P. Syndecan-1 interaction with the LG4/5 domain in laminin-332 is essential for keratinocyte migration. *J Cell Physiol* **214**, 238–249 (2008).

